# Shifting ecological niches of *Prunus* species under climate change

**DOI:** 10.1101/2025.01.17.633525

**Authors:** Chiara Vanalli, Marino Gatto, Renato Casagrandi, Daniele Bevacqua

**Affiliations:** Environmental Computational Science and Earth Observation Laboratory, École Polytechnique Fédérale de Lausanne (EPFL), 1950 Sion, Switzerland; Dipartimento di Elettronica, Informazione e Bioingegneria, Politecnico di Milano, 20133 Milano, Italy; INRAE, UR1115, Plantes et Systèmes de culture Horticoles (PSH), Site Agroparc, 84914 Avignon, France

**Keywords:** Temperature, Plant phenology, Dormancy break, Blooming, Fruit ripening, Adaptation

## Abstract

Climate change is strongly impacting agriculture, reducing crop production and shifting the geographic distribution of suitable areas for crop cultivation. To safeguard future global yield and feed a growing world population, the migration of crop production areas to new suitable sites represents a way to adapt to a changing climate. Here, we aim to identify the ecological niche of *Prunus* species, namely peach, plum, almond, apricot and sweet cherry, and examine their expected future shifts under climate change. For each *Prunus* species, we selected from the literature processed-based phenological models for dormancy break, blooming and fruit ripening, whose fulfillment determines whether an area is suitable for crop cultivation. Historically, the ecological niche of *Prunus* species spans mid-low European latitudes, while higher latitudes fail satisfying the forcing requirements for blooming and fruit ripening. In future, this constraint is expected to become less restrictive with a northwards expansion of the suitable areas. However, this will be contrasted by a contraction of the niche at low latitudes due to dormancy break failures. While bridging established mechanistic knowledge on the climatic effects of plant phenological traits with citizen science observations, our work brings new insights into how fruit crops will respond to global warming.

## Introduction

Climate change is modifying the global weather patterns, altering the frequency and intensity of extreme weather events and raising the average temperature across the globe^1^. Such changes are impacting the agricultural sector, with a reduced productivity growth and an increased vulnerability in global crop yields^2,3^, together with a shift of the geographic distribution of suitable areas for crop cultivation^4–8^. Estimates indicate an expected reduction of the global crop yield from 3% to 7% for each degree-Celsius of temperature increase^9–11^ and a reduced cropland suitability at lower latitudes^5,6^. Yet, the world population is growing and is expected to demand 70% more food by the middle of the XXI century^12^. In light of the described disparity between possible reduction in food production and population growth, the achievement of the United Nations “Zero Hunger” Sustainable Development Goal by 2030 is a challenging target^13^ and asks for well-designed adaptation strategies that can move agriculture towards a more productive and sustainable system.

Examining whether and how the suitable areas for crop cultivation will be shifted due to a changing environment can safeguard future global yield, guiding climate change adaptation through migration of crop production areas to new suitable sites^14–16^. This is particularly relevant for the cultivation of perennial crop whose lifespan covers multiple years and require an adequate agro-economic planning. The study of suitable geographic range for the cultivation of a specific crop dives into the theory of the ecological niche, i.e., the environmental conditions that a species requires to survive and maintain its populations^17^. Ecological niche modeling (or species distribution modeling) studies where a species can live and thrive in relation to environmental conditions. This is done either by examining the environmental characteristics of areas where the species is already known to exist or by incorporating detailed knowledge of how environmental factors influence the species’ traits, thereby affecting its survival^18,19^. Plant phenology, impacting key events in the plant annual cycle like dormancy, blooming and fruit ripening^20,21^, is recognized as one of the most important climate-dependent determinants that shape the ecological niche of a species^22–24^. Sifts in plant phenology are among the most significant documented responses to climate change^25–31^. Two prominent trends include earlier blooming times^26,28,32^ and shorter fruit development periods^33^. However, in the future, additional trends could emerge due to increasingly mild winter temperatures. Such conditions may prevent flowering in certain plant species if their chilling requirements are not met^34–38^. Failure in the fulfillment of the chilling requirements for dormancy break can disrupt the phenological phases necessary for growth and reproduction, compromising plant blooming, fruit development and indeed the crop yield^39–41^. Thus, suitable areas for crop cultivation require an environment characterized by favorable climatic conditions for the fulfillment of the different phenological requirements of the plant annual cycle: low temperatures are necessary to break bud dormancy, while higher temperatures are necessary to promote bud growth for blooming and fruit ripening. Conversely, locations where the phenological demands are failed to be met are excluded from the ecological niche of the considered crop. Therefore, an improved knowledge of the climatic requirements for the achievement of the different phases of the plant phenological cycle provides essential understanding to study the climate-driven shifts of the ecological niche and the agricultural suitability of crops.

Process-based phenological models that explicitly consider the different phases of the plant annual cycle are mainly split into one-phase models, which considered only the heat requirements^42,43^, and two-phase models, which instead assume a required amount of cold temperatures experienced before the heat requirement for blooming is achieved^44–46^. While the one-phase modeling framework could perform well in estimating plant phenology in past conditions, when the fulfillment of winter chilling needs was taken for granted, the two-phase models are preferred to project phenology for future climatic conditions^42,43^, given the limit that dormancy break failures could impose to crop cultivation^31^. After blooming, fruit growth and ripening are also influenced by climatic conditions. This process is commonly modeled through a Growing Degree Day (GDD) requirement, where enough GDD units need to be cumulated in order to achieve full ripening^38,47^. In addition to a well-established research area in phenological modeling, recent decades have seen a rapid growth in observational data gathered through citizen science platforms, providing valuable insights for modeling species distribution and environmental suitability^48,49^. However, approaches based on these large datasets often focus on empirical evidence, overlooking the underlying processes that drive the observed patterns^23^.

In the present work, we aim to identify the ecological niche of *Prunus* species, namely peach (*P. persica*), plum (*P. domestica*), almond (*P. amygdalus*), apricot (*P. armeniaca*) and sweet cherry (*P. avium*) and examine whether and how climate change is expected to reshape it. To do so: *i*) we performed a literature analysis on calibrated processed-based phenological models for dormancy break, blooming and fruit ripening; *ii*) we selected the most robust and comprehensive modeling frameworks and we implemented them to assess the historical ecological niche for the considered five species across the European territory; *iii*) we validated model predictions using presence observations in Europe provided by the Global Biodiversity Information Facility (GBIF)^50^; *iv*) we projected future shifts of the niche for each *Prunus* species across the XXI century under two different climate change scenarios, a more moderate scenario of sustainability, SSP1-2.6, and a more extreme climatic scenario of fossil-fuel-centered development, SSP5-5.8^1^. By integrating established mechanistic knowledge of the climatic effects on plant phenological traits with novel data sources, such as citizen science observations, our work offers new insights into how key species in the European agroeconomy, particularly fruit crops, will respond to global warming.

## Results

Findings from the performed literature review of available climate-driven process-based phenological models for *Prunus* species highlight the important role that temperature plays for the different phases of the plant phenological cycle and the variety of existing modeling frameworks (Table 1 and Additional Information). For the plant dormancy phase, cold requirements calculated through the Weinberger model (Chilling Hours)^51^ and the Utah model (Chilling Units)^52^ are usually outperformed by the Dynamic model (Chilling Portions)^53–55^, which considers a more complex chilling accumulation in a two-step process requiring both cold and mild temperatures. The dynamic model was used to model the climatic dependence of dormancy break for plum, almond and cherry, while the Chuine model^42,45^ was selected for peach and apricot (Table 1). For the remaining phenological phases of blooming and fruit ripening, there is an agreement across the examined frameworks that an accumulation of forcing units, either from a Growing Degree Day/Hour models (GDD, GDH) or from a sigmoidal function of temperature is required. Therefore, by bridging process-based modeling frameworks already available in literature, we were able to reconstruct the constrasting climatic dependence of the multiple phenologial phases and to map the phenological suitability of the European territory for the cultivation of peach (*P. persica*), plum (*P. domestica*), almond (*P. amygdalus*), apricot (*P. armeniaca*) and sweet cherry (*P. avium*).

**Table 1.** Selected phenological models for dormancy break, blooming and fruit ripening for the different *Prunus* species. CP=Chilling portions, GDH=Growing Degree Hours, GDD= Growing Degree Days; in italics the model inputs are reported (Tmean=mean temperature, Tmin=minimum temperature and Tmax=maximum temperature).

| Species | Dormancy break | Blooming | Ripening |
| --- | --- | --- | --- |
| Peach | Chuine model<sup>38</sup><br>*Daily Tmean* | Sigmoid model<sup>38</sup><br>*Daily Tmean* | GDD<sup>38</sup><br>*Daily Tmean* |
| Plum | CP<sup>62</sup><br>*Hourly Tmean (Daily Tmin, Tmax)* | GDH<sup>62</sup><br>*Hourly Tmean (Daily Tmin, Tmax)* | GDD<sup>66</sup><br>*Daily Tmean* |
| Almond | CP<sup>47,61</sup><br>*Hourly Tmean (Daily Tmin, Tmax)* | GDD<sup>47,61</sup><br>*Daily Tmean* | GDD<sup>47,61</sup><br>*Daily Tmean* |
| Apricot | Chuine model<sup>42</sup><br>*Daily Tmean* | Sigmoid model<sup>42</sup><br>*Daily Tmean* | GDD<sup>65</sup><br>*Daily Tmean* |
| Cherry | CP<sup>63</sup><br>*Hourly Tmean (Daily Tmin, Tmax)* | GDH<sup>63</sup><br>*Hourly Tmean (Daily Tmin, Tmax)* | GDH<sup>64</sup><br>*Hourly Tmean (Daily Tmin, Tmax)* |

The overlap between the presence data from GBIF of each *Prunus* species (Figure 1a-j) and the simulated historical ecological niche (Figure 1k-o) shows a good global recall metrics^56^, namely recall equals 88.36% for peach, 87.03% for plum, 73.14% for almond, 95.76% for apricot, 91.85% for cherry. Specifically, the ecological niche of most species spans low and mid European latitudes with the exception of mountainous areas, such as Alps, Carpathians and Pyrenees. The almond niche is that one that differentiates most from the others, with the simulated suitability for almonds located only in southern Europe and also confirmed by presence data. This peculiarity is explained by the phenological model with a higher requirement of forcing units, needed for almond growth and hull split^47^ compared to the other crops, which are stone fruits and present a lower forcing requirement. The selected models reproduce well the main production areas of central Europe, but fail to include some locations where presence data are instead reported (k-o). These regions of poor model performance mainly involve plums and cherries and are mainly located in the Alps and in center-northern UK and can be due to specific cultivars that are selected for those areas which are not well represented by the considered phenological models and/or to microclimatic conditions that are playing an important role in these locations.

**Figure 1.**
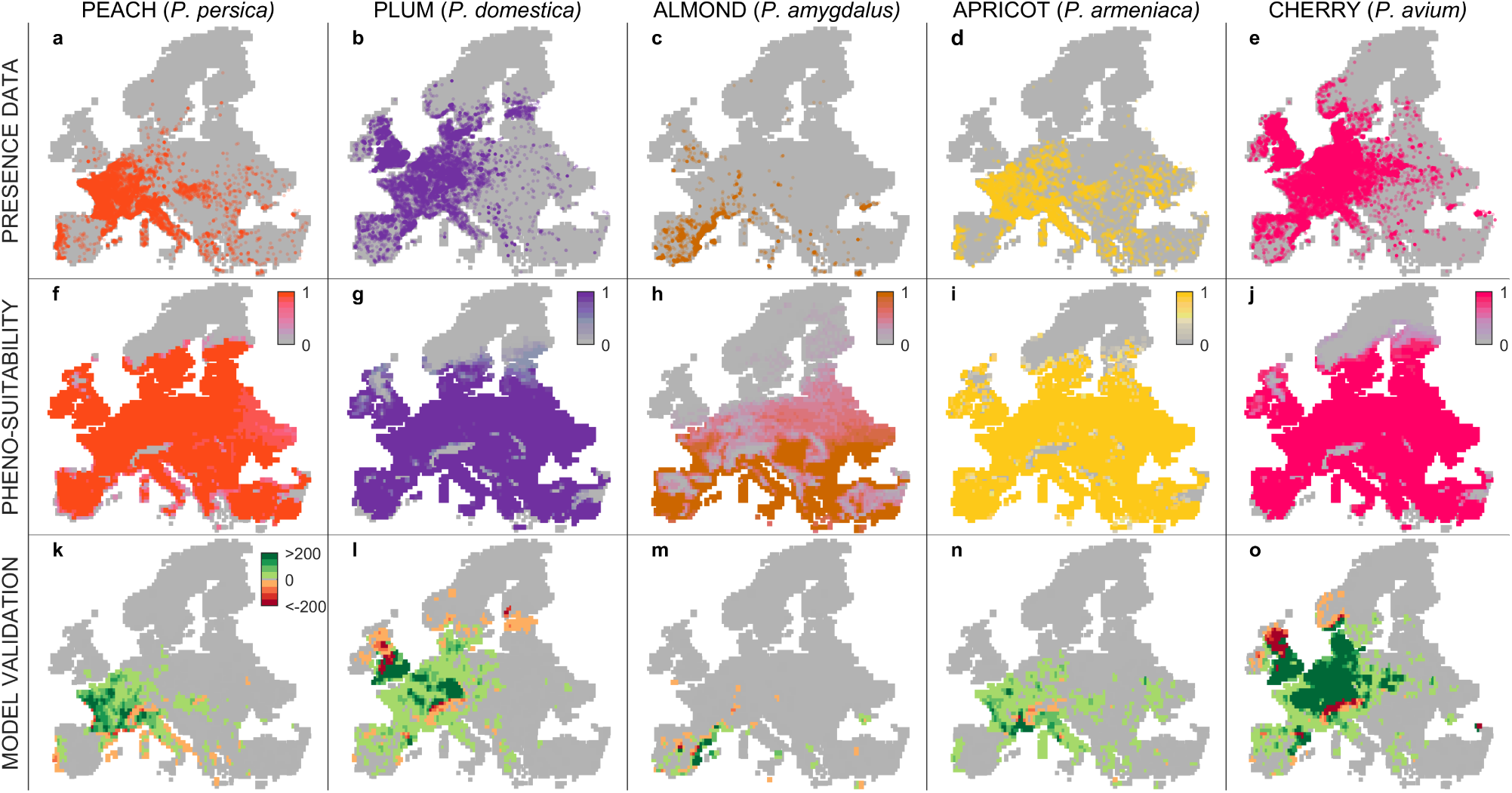
(a-e) Presence data and (f-j) simulated historical phenological suitability for peach (*Prunus persica*) in orange, plum (*Prunus domestica*) in purple, almond (*Prunus amygdalus*) in brown, apricot (*Prunus armeniaca*) in yellow and sweet cherry (*Prunus avium*) in magenta (suitability is rescaled from 0 to 1 according to the number of years over the 20 year-period for which the plant fulfills the phenological requirements); (k-o) spatial performance of model validation with the number of observed presences within a simulated suitable spatial cell (i.e. true positives) represented in green color scale, while the number of of observed presences within a simulated unsuitable spatial cell (i.e. false negatives) is plotted using a red color scale.

We projected the expected future changes in the ecological niche of the considered *Prunus* species at the end of the century under two different climate change scenarios, a sustainabile pathway (SSP1-2.6) and an unconstrained energy use pathway (SSP5-8.5)^1^. Under both scenarios, we expect an expansion of the suitable areas for *Prunus* cultivation at northern latitudes and in the mountain regions (Figure 2). This shift is explained by warmer temperatures that could allow for the fulfillment of forcing requirements for blooming and ripening. Conversely, in the Mediterranean basin a contraction of the suitable areas for *Prunus* cultivation is expected because of too mild winter temperatures that could impede the fulfillment of the chilling requirements for dormancy break (Figure 2). These changes are projected to be more consistent in the most extreme climate pathway (SSP5-8.5) compared to the more sustainable one (SSP1-2.6). Among the analyzed *Prunus* species, the almond niche is expected to expand the most under future climate change due to its high threshold of GDD requirements for ripening that, in the historical period, was fulfilled only in few Mediterranean areas. The suitability changes at the middle of the century are in line with those described for the end of the century, but they are less significant (Figure S1,S2).

**Figure 2.**
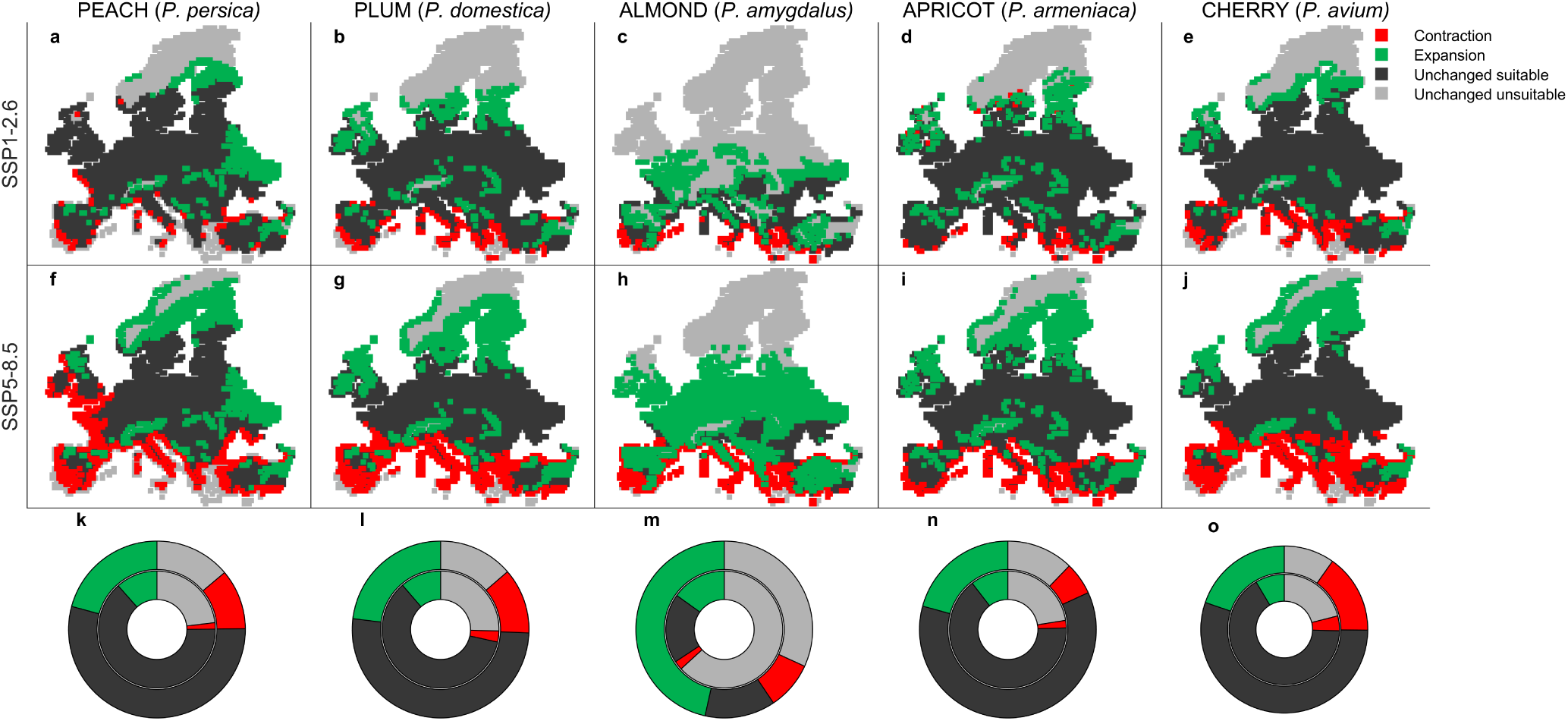
Future shifts of the ecological niche at the end of the XXI century (2081-2100), compared to the historical period (1995-2014) under a sustainability scenario of climate change SSP1-2.6 (a-e) and a fossil-fuel development scenario SSP5-8.5 (f-j). Red areas represent a future niche contraction, green areas a future expansion, light gray areas are unchanged unsuitable areas and dark gray areas represent no niche change compared to the historical period. (k-o) Composition of the European territory between areas of contraction, expansion and unchanged in the future under the SSP1-2.6 scenario (inner donuts) and under the SSP5-8.5 scenario (outer donuts), compared to the historical period.

Examining the latitudinal gradient of the suitable areas for *Prunus* cultivation, the historical niche for most species is located between 40 and 55 degrees of latitude except for almonds, whose niche is located at a lower latitudinal gradient, between 35 and 45 degrees (Figure Figure 3a-e). According to the phenological requirements, at the end of the century our findings indicate an average latitudinal shift of +1.31,+4.02 degrees, +0.832,+3.07 for peach, +1.34,+4.18 for plum, +2.18,+5.68 for almond, +0.818,+2.76 for apricot and +1.36, +4.43 for cherry. European historical unsuitable areas for *Prunus* cultivation are mainly characterized by a too strict climate that fails to satisfy the forcing requirements (Figure 3f-j, black lines). Those unsuitable locations are distributed not only at northern latitudes, but also where European mountain chains are located. On the one hand, warmer future climate could drastically decrease forcing failures at high latitudes, on the other hand, growing forcing failures are expected at lower latitudes in particular for the SSP 5-8.5 scenario. This opposite southern trend is probably caused by a very late fulfillment of the chilling requirement for dormancy break that does not give enough thermal time to the plant to achieve blooming and then to the fruit to develop and ripe. Failure areas for chilling requirements represent a less important constraint for the *Prunus* suitability in the historical period. However, too mild future winters are expected to increase chilling failures, with a contraction of the ecological niche at low latitudes (Figure 3f-j, gray lines). These latitudinal trends, albeit of a smaller magnitude, are similar to those projected for the middle of the century, when chilling failures at low latitudes are already showing to undermine the *Prunus* phenological suitability (Figure S1,S2). Overall, the decreased forcing failures in northern Europe will outbalance the increase in chilling failures in southern Europe with a net expansion of the niche of *Prunus* (peach +12,+14%, plum +13,+15%, almond +58,+174%, apricot +12,+20%, cherry +3,+6%, Figure 3k). Yet, to benefit from these new suitable areas, proper planning of migration of cultivation areas is required that needs to take into consideration not only the climatic suitability for plant phenology, but also agro-economic and land use characteristics.

**Figure 3.**
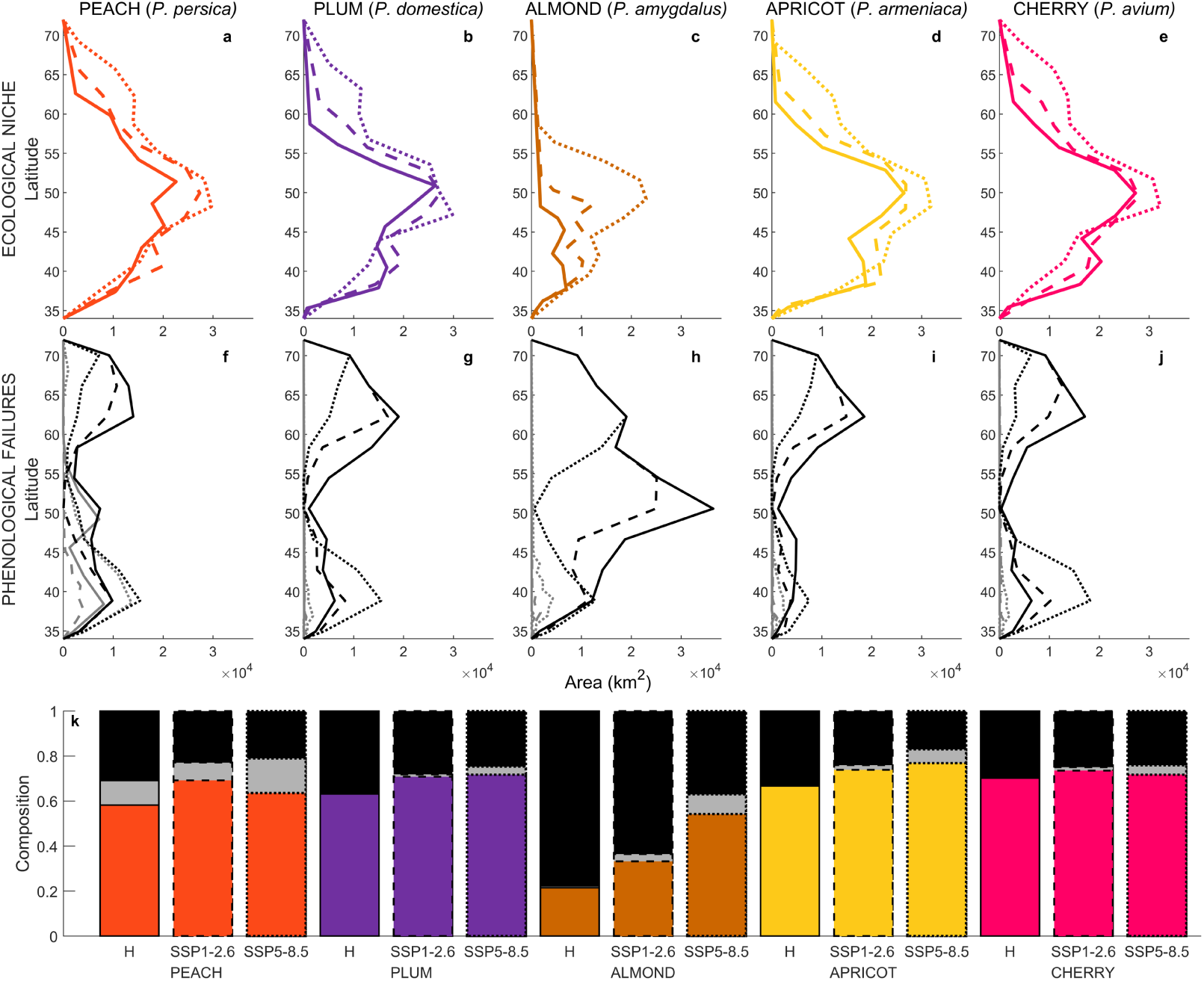
Latitudinal gradient of *Prunus* (a-e) ecological niche and (f-j) phenological failures (in gray chilling failures and in black forcing failures) in the historical period (1995-2014, solid line), at the end of the century under SSP1-2.6 scenario (2081-2100, dashed line), and at the end of the century under SSP5-8.5 scenario (2081-2100, dotted line). (k) Barplot represents the composition of the European territory between suitable areas (colored in orange for peach, purple for plum, brown for almond, yellow for apricot and magenta for cherry) in gray chilling failures and in black forcing failures, during the historical period (H, solid edge) and at the end of the century under the two scenarios (SSP1-2.6 scenario dashed bar edge, SSP5-8.5 scenario dotted bar edge).

## Discussion

To face the challenges that climate change poses to agriculture, the examination of whether and how the suitability of crop cultivation is impacted by global warming is required to adequately adapt and guarantee a sufficient future yield. In this study, we focused on five *Prunus* species and evaluated their current and future niches in Europe based on species-specific phenological requirements for dormancy break, blooming, and fruit ripening, recovered from a literature analysis. Historically, the *Prunus* niche estimated from phenological suitability overlaps quite well with the presence data retrieved from GBIF^50^. The main historical limitation for *Prunus* cultivation for the European territory is represented by the forcing requirements for blooming and fruit ripening that cannot be satisfied in cold environments at central-high latitudes. Conversely, at the end of the XXI century, this constraint is expected to become less restrictive with a northwards expansion of the suitability niche of +1.31,+4.02 latitudinal degrees on average. This positive expansion will be contrasted by the increase in dormancy break failures in the Mediterranean basin due to too mild winters that would not allow the fulfillment of the chilling requirements. The described general trends of crop suitability and phenology that we highlighted in the current study for the *Prunus* genus in response to the predicted increase in temperature have been predicted for Europe for other economically important fruit crops, such as grapes^57^ and olives^58^. Altogether, this represents an alarming trend for the Mediterranean basin, where most of the production areas are currently located.

In our work, we leveraged citizen science observations on presences of *Prunus* species across Europe, which generally represent an enormous resource for scientific research and in particular for planetary-scale challenges such as global warming^59^. However, the use of citizen science data is related to several biases^60^: *i*) absences are usually outweighed by presences and have not been considered in the current work, *ii*) spatial and temporal distribution of records depend on many factors including the popularity of these platforms, *iii*) observations can be related to ornamental cultivars of the considered species rather than actual agricultural crops. However, these biases play a minor role in research studies, like ours, where citizen science data are used to validate existing models which have been previously calibrated using field and/or experimental phenological observations. Overall, the inclusion plant phenological stages in addition to the presence records in citizen science data would greatly benefit scientific research on climatic impacts on plant phenology and distribution.

Our study is based on process-based phenological modeling frameworks from the literature^38,42,47,61–66^ that provided crucial knowledge on the climatic response of the different plant phenological stages. Plant phenology and its climatic dynamics are rarely addressed in ecological niche modeling, which usually disregards the link between specific phenological stages and environmental variables, potentially producing unreliable future projections under rapid climatic changes^67,68^. In addition, the existing phenological works usually focus on one specific phenological stage of the plant annual cycle (but see^38,47^). The evaluation of the entire annual plant phenological cycle is instead needed to achieve a more complete understanding of climate change impacts across the multiple plant climatic requirements. In our phenological framework, for each species and phenological stage, we selected the highest scoring model based on its robustness and completness. A different approach could have been to ensemble the multiple models available in the literature^69,70^. An additional level of model complexity is represented by cultivar-specific parametrization of the selected models. In absence of cultivar-level presence data, we decided to choose the most permissive parameters, assuming that a spatial cell is suitable for a specific crop if at least one cultivar can achieve fruit ripening. Differences in plant phenological requirements between different cultivars represent one potential adaptation strategy thanks to the selection of specific cultivars with a low chilling requirement for dormancy break^71^. Ultimately, the present work is based on the phenological thermal requirements of *Prunus* species and maps the potential ecological niche of the examined species, neglecting additional biotic interactions that could play a relevant role in the realized niche^17^. For instance, pollinators are essential to complete the plant phenological cycle and are undermined by climate change^72^. To effectively design crop migration strategies to adapt to future climatic conditions, our considerations based on phenological climatic suitability need to be paired with targeted studies on economic feasibility and territorial characteristics.

To tackle the great challenges that climate change is posing to agriculture and food security, our study can be used to minimize the negative impacts of the changing climate, with an optimal migration of crop cultivation areas towards new suitable areas where the plant phenological requirements will still be fulfilled in the future.

## Methods

### Phenological models of *Prunus* species

We performed a literature analysis following the Preferred Reporting Items for Systematic reviews and Meta-Analyses (PRISMA) criteria to identify mechanistic phenological models of *Prunus* species available in the literature.

Search words: ((“Prunus”) AND (“persica” OR “domestica” OR “armeniaca” OR “dulcis” OR “avium”)) OR (“peach” OR “apricot” OR “plum” OR “almond” OR “cherry”) AND (“Phenology” OR “Blooming” OR “Flowering” OR “Chilling” OR “Ripening” OR “Fruit growth”) AND (“Temperature” OR “Climate” OR “Thermal” OR “Heat”) AND (“Model*“ OR “Simulat*“OR “Predict*“). Done on July 5^th^ 2024.

From more than 500 records identified, 95 were selected for full-text analysis (Peach (18), Plum (6), Almond (18), Apricot (24), Cherry (29), see Table S1). For plum (*P. domestica*), we also considered phenological models of *Prunus salicina* commonly called Japanese plum, which is commonly cultivated in Europe. For each species, we scored each model according to the following criteria:

- if model considers both dormancy and forcing=+1
- if data on dormancy break time are availble in addition to blloming=+1
- if blooming and ripening time are available from the same dataset=+1
- if phenological model is a result from literature review=+1
- if phenological data location used for model calibration is within Europe=+1
- if phenological data used for model calibration cover a time period of more than 15 years=+1
- if number of considered cultivars is more than 5=+1

We normalized the blooming and ripening score to 1 and we selected the phenological model with the highest score for each species. If multiple models present the same scores, models that compare different phenological modeling frameworks and/or include in their datasets multiple cultivars are preferred (Tables S2–S6). The selected phenological models with the relative temperature inputs are reported in Table 1.

### Occurrences of *Prunus* species for validation

We used presence-only data of *Prunus* species recorded from 1995 to 2024 from the Global Biodiversity Information Facility (GBIF) to validate the simulations of the selected phenological model across the European territory. For each species, as global metric of model performance we computed the model recall^56^:

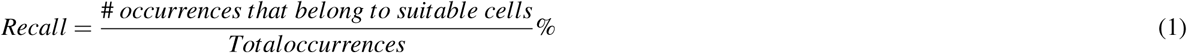

A spatial cell is considered suitable if the phenological requirements for dormancy break, blooming and ripening are satisfied for at least the 90% of the considered time periods. Among the unsuitable cells, we defined a chilling failure when the cold requirement for dormancy break are not satisfied, while a forcing failure when the heat requirements for blooming and/or for ripening are not met.

### Historical and future climatic data

We retrieved historical daily average, maximum and minimum air temperature from the Coupled Model Intercomparison Project Phase 6 (CMIP6, CNRM-CM6-1-HR model) at a spatial resolution of 0̃.5×0.5 degrees for the European territory (3,872 total cells)^73^. We analyzed a more sustainable Shared Socioeconomic Pathways (SSPs), SSP1-2.6, and a fossil-fuel oriented scenario, SSP5-8.5, with the same temporal and spatial resolution^1^. We evaluated the climatic data in the historical period, from 1994 to 2014, and in the future, under the two considered scenarios, at the middle of the century. from 2041 to 2060, and at the end of the century, from 2080 to 2100. We used the methods described in Lopez-Bernal et al. (2018)^74^ to compute the hourly temperatures from daily values, which are the climatic inputs for the phenological models that runs with hourly time step.

## Acknowledgements

We would like to acknowledge Dr. Devis Tuia for his valuable support.

## Author contributions statement

CV and DB developed the original idea. CV performed the literature analysis, modeling work, result visualization and wrote the first draft of the manuscript. DB, MG and RC guided the modeling development advised on the analysis and results interpretation. All authors reviewed the manuscript.

## Data availability

The datasets used in the current study are publicly available on GBIF (https://www.gbif.org/) and on Copernicus (https://cds.climate.copernicus. and can also be requested contacting the corresponding author.

## Additional information

**Table S1.** Literature review of process-based pehnological models of *Prunus* species: reference ID, authors, year of publication and journal.

| Ref ID | Authors | Year | Journal |
| --- | --- | --- | --- |
| 1 | Pantelidis et al. | 2023 | Scientia Horticulturae |
| 2 | Pantelidis et al. | 2022 | Acta Horticulturae |
| 3 | Cifuentes-Carvajal et al. | 2023 | Agronomy |
| 4 | Drogoudi et al. | 2023 | Plants |
| 5 | Chuine et al. | 2016 | Global change biology |
| 6 | Atagul et al. | 2022 | Agronomy |
| 7 | Schwartz et al. | 1997 | HortScience |
| 8 | Marra et al. | 2002 | Acta Horticulturae |
| 9 | Vanalli et al. | 2021 | Agricultural and Forest Meteorology |
| 10 | Kwon et al. | 2020 | Agricultural and Forest Meteorology |
| 11 | Maulión et al. | 2014 | Scientia Horticulturae |
| 12 | Lu et al. | 2012 | Acta Horticulturae |
| 13 | Miranda et al. | 2013 | Agricultural and Forest Meteorology |
| 14 | Kamyun et al. | 2014 | Acta Horticulturae |
| 15 | Huret et al. | 2017 | International Journal of Climatology |
| 16 | Bonhomme et al. | 2010 | Acta Horticulturae |
| 17 | Kwon et al. | 2020 | Acta Horticulturae |
| 18 | Borgini et al. | 2024 | Scientia Horticulturae |
| 19 | Guerrero et al. | 2024 | Frontiers in Plant Science |
| 20 | Hamdani et al. | 2024 | Theoretical and Applied Climatology |
| 21 | Guerrero et al. | 2022 | Acta Horticulturae |
| 22 | Fraga et al. | 2021 | Frontiers in Plant Science |
| 23 | Borgini et al. | 2024 | Scientia Horticulturae |
| 24 | Orjuela-Angulo et al. | 2022 | Revista Colombiana de Ciencias Hortícolas |
| 25 | Kumawat et al. | 2023 | Erwerbs-Obstbau |
| 26 | Orozco et al. | 2024 | Scientific Reports |
| 27 | Naseri et al. | 2023 | Theoretical and Applied Climatology |
| 28 | Degrandi-Hoffman et al. | 1996 | Journal of Applied Ecology |
| 29 | Alonso et al. | 2005 | Journal of the American Society for Horticultural Science |
| 30 | Ramírez et al. | 2010 | Acta Horticulturae |
| 31 | Freitaset al. | 2023 | Climatic Change |
| 32 | Lorite et al. | 2020 | Agricultural and Forest Meteorology |
| 33 | Ruml et al. | 2020 | Acta Horticulturae |
| 34 | Benmoussa et al. | 2017 | Agricultural and Forest Meteorology |
| 35 | Prudencio et al. | 2019 | Acta Horticulturae |
| 36 | Gaeta et al. | 2018 | European Journal of Agronomy |
| 37 | Prudencio et al. | 2018 | Scientia Horticulturae |
| 38 | Parker et al. | 2018 | Climatic Change |
| 39 | Parker et al. | 2017 | International Journal of Biometeorology |
| 40 | Prudencio et al. | 2018 | Acta Horticulturae |
| 41 | Gaeta et al. | 2018 | Acta Horticulturae |
| 42 | Santoset al. | 2017 | Climatic Change |
| 43 | Pantelidis et al. | 2023 | Scientia Horticulturae |
| 44 | Korsakova et al. | 2023 | Inventions |
| 45 | Vávraet al. | 2022 | Acta Horticulturae |
| 46 | Mesterházy et al. | 2022 | Diversity |
| 47 | Chuine et al. | 2016 | Global change biology |
| 48 | Andreini et al. | 2014 | Agricultural and Forest Meteorology |
| 49 | García et al. | 1999 | Acta Horticulturae |
| 50 | Legave et al. | 2006 | Acta Horticulturae |
| 51 | Valentini et al. | 2004 | Acta Horticulturae |
| 52 | Legave et al. | 2010 | Acta Horticulturae |
| 53 | Hangyu et al. | 2021 | Proceedings-ISCIPT 2021 |
| 54 | Ruiz et al. | 2007 | Environmental and Experimental Botany |
| 55 | Ruml et al. | 2012 | Acta Horticulturae |
| 56 | Campoyet al. | 2012 | European Journal of Agronomy |
| 57 | Bartolini et al. | 2019 | Scientia Horticulturae |
| 58 | Černá et al. | 2012 | Acta Universitatis Agriculturae et Silviculturae Mendelianae Brunensis |
| 59 | Guo et al. | 2015 | Scientia Horticulturae |
| 60 | Gao et al. | 2012 | HortScience |
| 61 | Bartolini et al. | 2020 | Acta Horticulturae |
| 62 | Zhuang et al. | 2016 | European Journal of Agronomy |
| 63 | Ruml et al. | 2018 | Acta Horticulturae |
| 64 | Kwon et al. | 2020 | Acta Horticulturae |
| 65 | Borgini et al. | 2024 | Scientia Horticulturae |
| 66 | Ruml et al. | 2010 | International journal of biometeorology |
| 67 | Ortega-Farias et al. | 2023 | Proceedings-IEEE CHILEAN |
| 68 | Fadón et al. | 2023 | Agricultural and Forest Meteorology |
| 69 | Fadón et al. | 2022 | Acta Horticulturae |
| 70 | Chmielewski et al. | 2023 | Horticulturae |
| 71 | Litschmann et al. | 2022 | Acta Horticulturae |
| 72 | Imperiale et al. | 2022 | Acta Horticulturae |
| 73 | Chmielewski et al. | 2022 | Plants |
| 74 | Fadón et al. | 2023 | European Journal of Agronomy |
| 75 | Campillay-Llanos et al. | 2024 | Scientia Horticulturae |
| 76 | Sugiura et al. | 2006 | Acta Horticulturae |
| 77 | Fadón et al. | 2021 | Agricultural and Forest Meteorology |
| 78 | Chozinski et al. | 2008 | Acta Horticulturae |
| 79 | Hochmaier | 2014 | Acta Horticulturae |
| 80 | Albuquerque et al. | 2008 | Environmental and Experimental Botany |
| 81 | Darbyshire et al. | 2020 | Agricultural and Forest Meteorology |
| 82 | Luedeling et al. | 2013 | International Journal of Biometeorology |
| 83 | Wenden et al. | 2018 | Acta Horticulturae |
| 84 | Guak et al. | 2013 | Horticulture Environment and Biotechnology |
| 85 | Persely et al. | 2014 | Acta Horticulturae |
| 86 | Drogoudi et al. | 2020 | Euro-Mediterranean Journal for Environmental Integration |
| 87 | Kaufmann et al. | 2017 | Regional Environmental Change |
| 88 | Palasciano et al. | 2017 | Acta Horticulturae |
| 89 | Chung et al. | 2016 | Asia-Pacific Journal of Atmospheric Sciences |
| 90 | Marra et al. | 2017 | Acta Horticulturae |
| 91 | Chmielewski et al. | 2016 | Agricultural and Forest Meteorology |
| 92 | Hur et al. | 2017 | International Journal of Climatology |
| 93 | Luedeling et al. | 2013 | Agricultural and Forest Meteorology |
| 94 | Fadón et al. | 2021 | Acta Horticulturae |
| 95 | Schiffers et al. | 2021 | Acta Horticulturae |

**Table S2.**
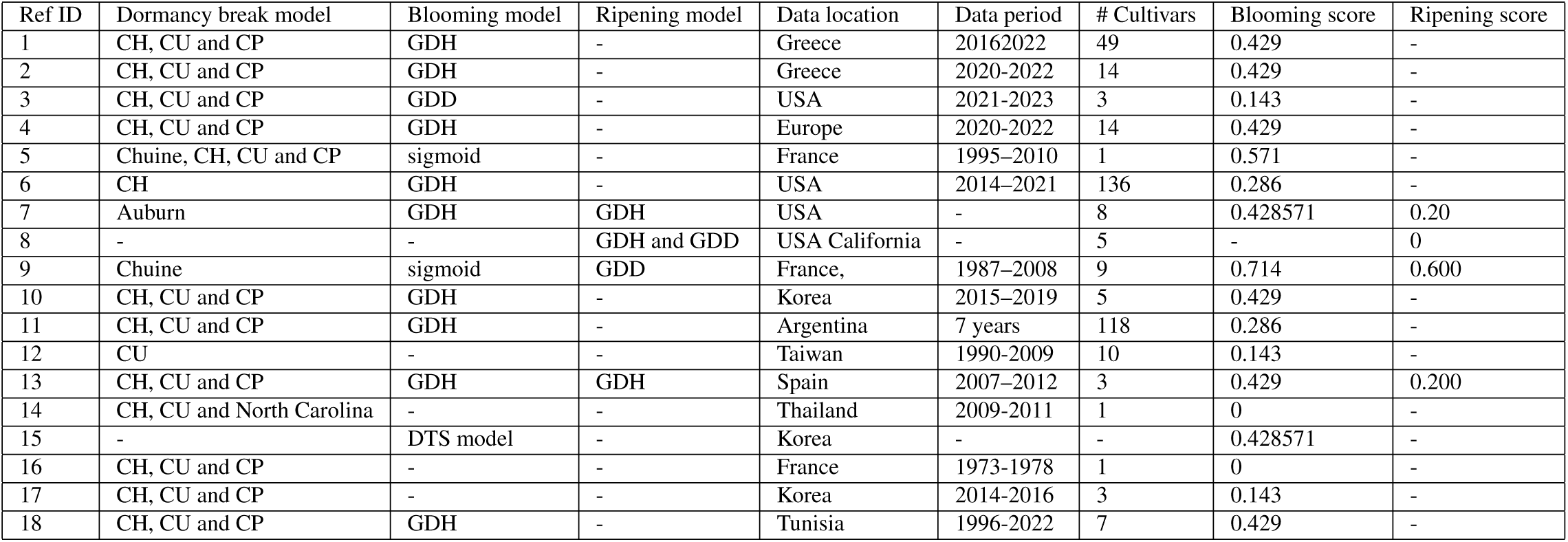
Literature review of peach phenological models together with model scoring for blooming and ripening. CH=chilling hours, CU=chilling unit, CP=chilling portions, NC= North Carolina model, GDD=Growing Degree Days, GDH=Growing Degree Hours.

| Ref ID | Dormancy break model | Blooming model | Ripening model | Data location | Data period | # Cultivars | Blooming score | Ripening score |
| --- | --- | --- | --- | --- | --- | --- | --- | --- |
| 1 | CH, CU and CP | GDH | - | Greece | 2016-2022 | 49 | 0.429 | - |
| 2 | CH, CU and CP | GDH | - | Greece | 2020-2022 | 14 | 0.429 | - |
| 3 | CH, CU and CP | GDD | - | USA | 2021-2023 | 3 | 0.143 | - |
| 4 | CH, CU and CP | GDH | - | Europe | 2020-2022 | 14 | 0.429 | - |
| 5 | Chuine, CH, CU and CP | sigmoid | - | France | 1995-2010 | 1 | 0.571 | - |
| 6 | CH | GDH | - | USA | 2014-2021 | 136 | 0.286 | - |
| 7 | Auburn | GDH | GDH | USA | - | 8 | 0.428571 | 0.20 |
| 8 | - | - | GDH and GDD | USA California | - | 5 | - | 0 |
| 9 | Chuine | sigmoid | GDD | France, | 1987-2008 | 9 | 0.714 | 0.600 |
| 10 | CH, CU and CP | GDH | - | Korea | 2015-2019 | 5 | 0.429 | - |
| 11 | CH, CU and CP | GDH | - | Argentina | 7 years | 118 | 0.286 | - |
| 12 | CU | - | - | Taiwan | 1990-2009 | 10 | 0.143 | - |
| 13 | CH, CU and CP | GDH | GDH | Spain | 2007-2012 | 3 | 0.429 | 0.200 |
| 14 | CH, CU and North Carolina | - | - | Thailand | 2009-2011 | 1 | 0 | - |
| 15 | - | DTS model | - | Korea | - | - | 0.428571 | - |
| 16 | CH, CU and CP | - | - | France | 1973-1978 | 1 | 0 | - |
| 17 | CH, CU and CP | - | - | Korea | 2014-2016 | 3 | 0.143 | - |
| 18 | CH, CU and CP | GDH | - | Tunisia | 1996-2022 | 7 | 0.429 | - |

**Table S3.** Literature review of plum phenological models together with model scoring for blooming and ripening. See Table 1 caption for abbreviations.

| Ref ID | Dormancy break model | Blooming model | Ripening model | Data location | Data period | # Cultivars | Blooming score | Ripening score |
| --- | --- | --- | --- | --- | --- | --- | --- | --- |
| 19 | CH, CU and CP | GDH | - | Spain | 1973-2023 | 21 | 0.714 | - |
| 20 | CH, CU and CP | GDH | - | Marocco | 2021-2022 | 28 | 0.429 | - |
| 21 | CH, CU and CP | GDH | - | Spain | 2018-2020 | 8 | 0.571 | - |
| 22 | CP | GDH | - | Portugal | - | 1 | 0.429 | - |
| 23 | CP | GDH | - | Tunisia | 1996- 2022 | 11 | 0.286 | - |
| 24 | - | - | GDD | Colombia | 2015-2016 | 1 | - | 0 |

**Table S4.** Literature review of almond phenological models together with model scoring for blooming and ripening. See Table 1 caption for abbreviations.

| Ref ID | Dormancy break model | Blooming model | Ripening model | Data location | Data period | # Cultivars | Blooming score | Ripening score |
| --- | --- | --- | --- | --- | --- | --- | --- | --- |
| 25 | - | GDD | - | Kashmir | 2012–2018 | 8 | 0.143 | - |
| 26 | - | Carbohydrate-temperature | - | California | 1982–2022 | - | 0.143 | - |
| 27 | CH | GDH | - | Iran | 2017 | 3 | 0.286 | - |
| 28 | - | GDD | - | California | 1987-1990 | 5 | 0 | - |
| 29 | chilling units | GDH | - | World | 1994-2004 | 44 | 0.286 | - |
| 30 | CH, CU, CP, NC, Florida | GDH | - | Chile | 2003-2005 | 3 | 0.143 | - |
| 31 | CH and CP | GDH and GDD | - | Portugal | - | 9 | 0.429 | - |
| 32 | CP | GDH | - | Southern Europe | 2004-2015 | 15 | 0.429 | - |
| 33 | CU | GDH | - | Serbia | 2011-2014 | 6 | 0.286 | - |
| 34 | CH, CU and CP | GDH | - | Tunisia | 1981-2014 | 37 | 0.429 | - |
| 35 | CU and CP | GDH | - | Spain | 2014-2016 | 1 | 0.286 | - |
| 36 | CU and CP | GDH | - | Italy | 1982–1992 | 7 | 0.429 | - |
| 37 | CU and CP | GDH | - | Spain | 2015-2017 | 3 | 0.286 | - |
| 38 | CP | GDH | GDD | USA | 1971–2000 | - | 0.571 | 0.600 |
| 39 | CP | GDD | GDD | USA | 1971–2000 | - | 0.571 | 0.600 |
| 40 | CU and CP | GDH | - | Spain | 2014-2016 | 4 | 0.429 | - |
| 41 | CP | GDH | - | Italy | 1991-1993, 2002 | 5 | 0.286 | - |
| 42 | CP | GDH | - | Portugal | - | - | 0.286 | - |

**Table S5.** Literature review of apricot phenological models together with model scoring for blooming and ripening. See Table 1 caption for abbreviations.

| Ref ID | Dormancy break model | Blooming model | Ripening model | Data location | Data period | # Cultivars | Blooming score | Ripening score |
| --- | --- | --- | --- | --- | --- | --- | --- | --- |
| 43 | CH, CU and CP | GDH | - | Greece | 2016-2022 | 49 | 0.429 | - |
| 44 | Sigmoid, triangular | GDD | - | Russia | 1995-2022 | 15 | 0.571 | - |
| 45 | - | GDH and GDD | - | Czech Republic | 1998-2018 | - | 0.286 | - |
| 46 | Chuine | sigmoid | - | Hungary | 1994-2020 | 3 | 0.429 | - |
| 47 | Chuine | sigmoid | - | France | 1987–2008 | 1 | 0.571 | - |
| 48 | Sigmoid, CU, GDD, Bidabè | - | - | France, Italy, Spain | 1988-2009 | 23 | 0.429 | - |
| 49 | CU | GDH | - | Italy and Spain | 1991-1994 | - | 0.429 | - |
| 50 | Bidabè | GDH | - | France | 1998-2005 | 5 | 0.429 | - |
| 51 | Utah, North Carolina | GDH and GDD | - | Italy | 1999-2002 | 5 | 0.286 | - |
| 52 | CH, CU and Bidabé | - | - | France | 1997-2007 | 1 | 0.143 | - |
| 53 | - | GDD | - | China | 1980-2020 | - | 0.286 | - |
| 54 | CH, CU and CP | GDH | - | Spain | 2003-2005 | 10 | 0.286 | - |
| 55 | - | GDD | - | Serbia | 1995-2004 | 42 | 0.143 | - |
| 56 | CH, CU and CP | GDH | - | Spain, South Africa | 2006-2009 | 14 | 0.143 | - |
| 57 | CH | - | - | Italy | 1973–2016 | 40 | 0.429 | - |
| 58 | - | sine wave | - | Czech Republic | 1927–1960 | 1 | 0.429 | - |
| 59 | CH and CP | GDH | - | China | 1963–2010 | - | 0.286 | - |
| 60 | CH, CU and CP | GDH | - | China | 2009-2012 | 6 | 0.286 | - |
| 61 | CU | - | - | Italy | 1991-2016 | 2 | 0.429 | - |
| 62 | CP | GDH | - | China | 2009–2011 | 69 | 0.286 | - |
| 63 | CU | GDH | - | Serbia | 2011-2013 | 10 | 0.429 | - |
| 64 | CH, CU and CP | GDH | - | Korea | 1919-2019 | 2 | 0.429 | - |
| 65 | CH, CU and CP | GDH | - | Tunisia | 1996-2022 | 5 | 0.286 | - |
| 66 | - | - | GDD | Serbia | 1995-2004 | 10 | - | 0.200 |

**Table S6.** Literature review of cherry phenological models together with model scoring for blooming and ripening. See Table 1 caption for abbreviations.

| Ref ID | Dormancy break model | Blooming model | Ripening model | Data location | Data period | # Cultivars | Blooming score | Ripening score |
| --- | --- | --- | --- | --- | --- | --- | --- | --- |
| 67 | - | exponential of GDD | - | Chile | 2017-2021 | 8 | 0.143 | - |
| 68 | CP | GDH | - | Germany, Spain | 1978-2020 | 3 | 0.429 | - |
| 69 | CP | GDH | - | Spain | - | 12 | 0.429 | - |
| 70 | CP | GDH | - | Germany | 2011-2023 | 1 | 0.429 | - |
| 71 | CH, CU and CP | - | - | Czech Republic | 2017-2021 | 40 | 0.286 | - |
| 72 | CU and CP | GDH | - | Italy | 2019 | 6 | 0.429 | - |
| 73 | CP | GDH | - | - | 2011-2019 | 1 | 0.286 | - |
| 74 | CH, CU and CP | GDH | - | Spain | 1991-2022 | 7 | 0.571 | - |
| 75 | - | GDD | - | Chile | 2017-2021 | 7 | 0.143 | - |
| 76 | CH | - | - | - | 1995-1998 | 1 | 0 | - |
| 77 | CP | GDH | - | Spain | 22 years | 20 | 0.571 | - |
| 78 | - | - | GDH | USA | 2003-2004 | 7 | - | 0.200 |
| 79 | CH | GDH | - | Argentina | 2002-2007 | 4 | 0.286 | - |
| 80 | CH, CU and CP | GDH | - | Spain | 2004-2006 | 7 | 0.429 | - |
| 81 | sequential and overlap CP | GDH | - | Australia | 1992-2015 | 3 | 0.286 | - |
| 82 | CH, CU and CP | GDH | - | Germany | 1984-2008 | 1 | 0.286 | - |
| 83 | CP | Pope overlap | - | Europe | 1970-2018 | 1 | 0.429 | - |
| 84 | CH and CU | - | - | Canada | 2009-2010 | 1 | 0 | - |
| 85 | - | GDD | - | Hungary | 1984-1991 | 3 | 0.143 | - |
| 86 | CH, CU and CP | - | - | Germany, Greece | 1984-2019 | 4 | 0.286 | - |
| 87 | CH, CU and CP | - | - | Germany | 2012-2014 | 3 | 0.143 | - |
| 88 | CH, CU and CP | - | - | Italy | - | 10 | 0.286 | - |
| 89 | chilling days, anti-chilling days | - | - | South Korea | 1981-2010 | - | 0.143 | - |
| 90 | CH and CU | GDH | - | Italy | 1994-2013 | 4 | 0.143 | - |
| 91 | CP | GDD | - | Germany | 2001-2015 | 3 | 0.429 | - |
| 92 | - | DTS model | - | Korea | - | - | 0.429 | - |
| 93 | CP | GDH | - | Germany | 1984-2008 | 1 | 0.429 | - |
| 94 | CP | GDH | - | Germany | - | 1 | 0.286 | - |
| 95 | CP | sigmoidal | - | Germany | 1984-2018 | 1 | 0.429 | - |

**Figure S1.**
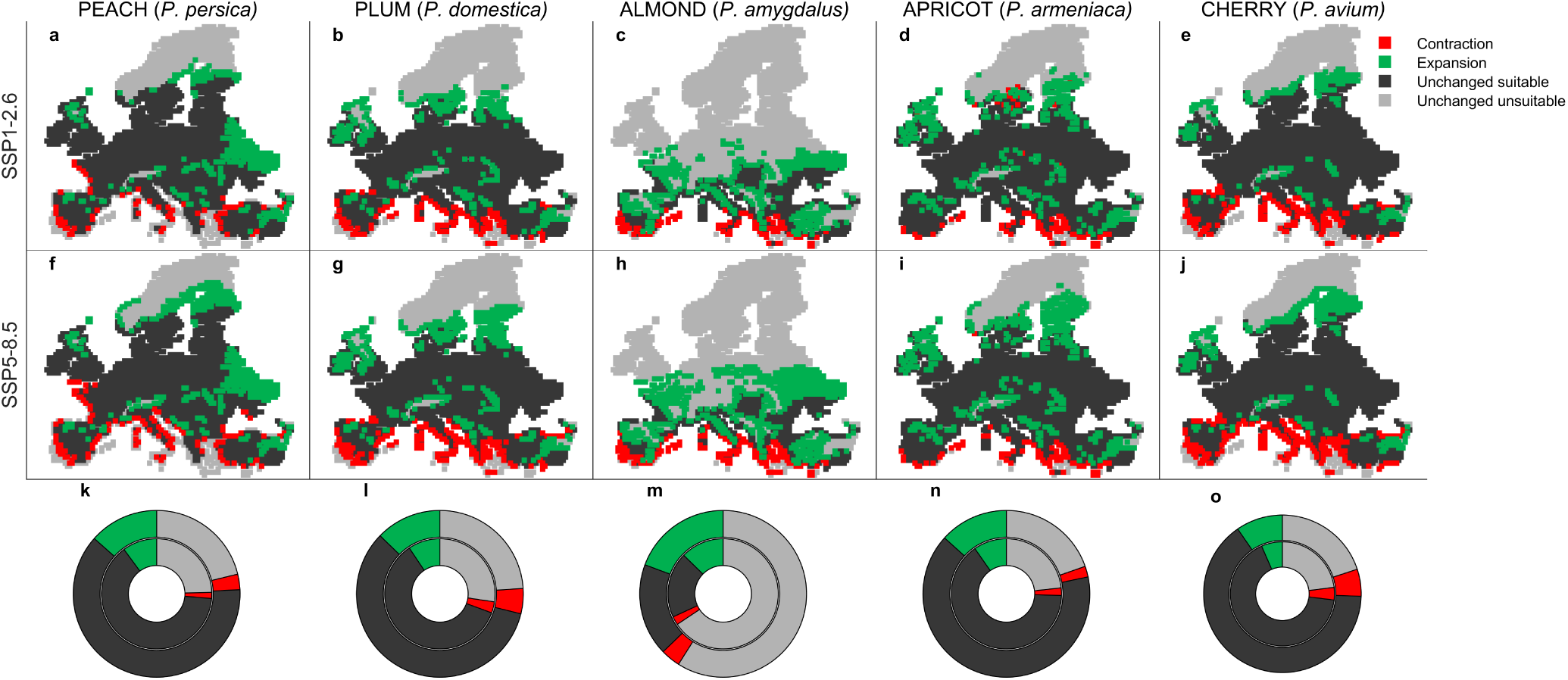
Future shifts of the ecological niche at the middle of the XXI century (2041-2060), compared to the historical period (1995-2014) under a sustainability scenario of climate change SSP1-2.6 (a-e) and a fossil-fuel development scenario SSP5-8.5 (f-j). Red areas represent a future niche contraction, green areas a future expansion, light gray areas are unchanged unsuitable areas and dark gray areas represent no niche change compared to the historical period. (k-o) Composition of the European territory between areas of contraction, expansion and unchanged in the future under the SSP1-2.6 scenario (inner donuts) and under the SSP5-8.5 scenario (outer donuts), compared to the historical period.

**Figure S2.**
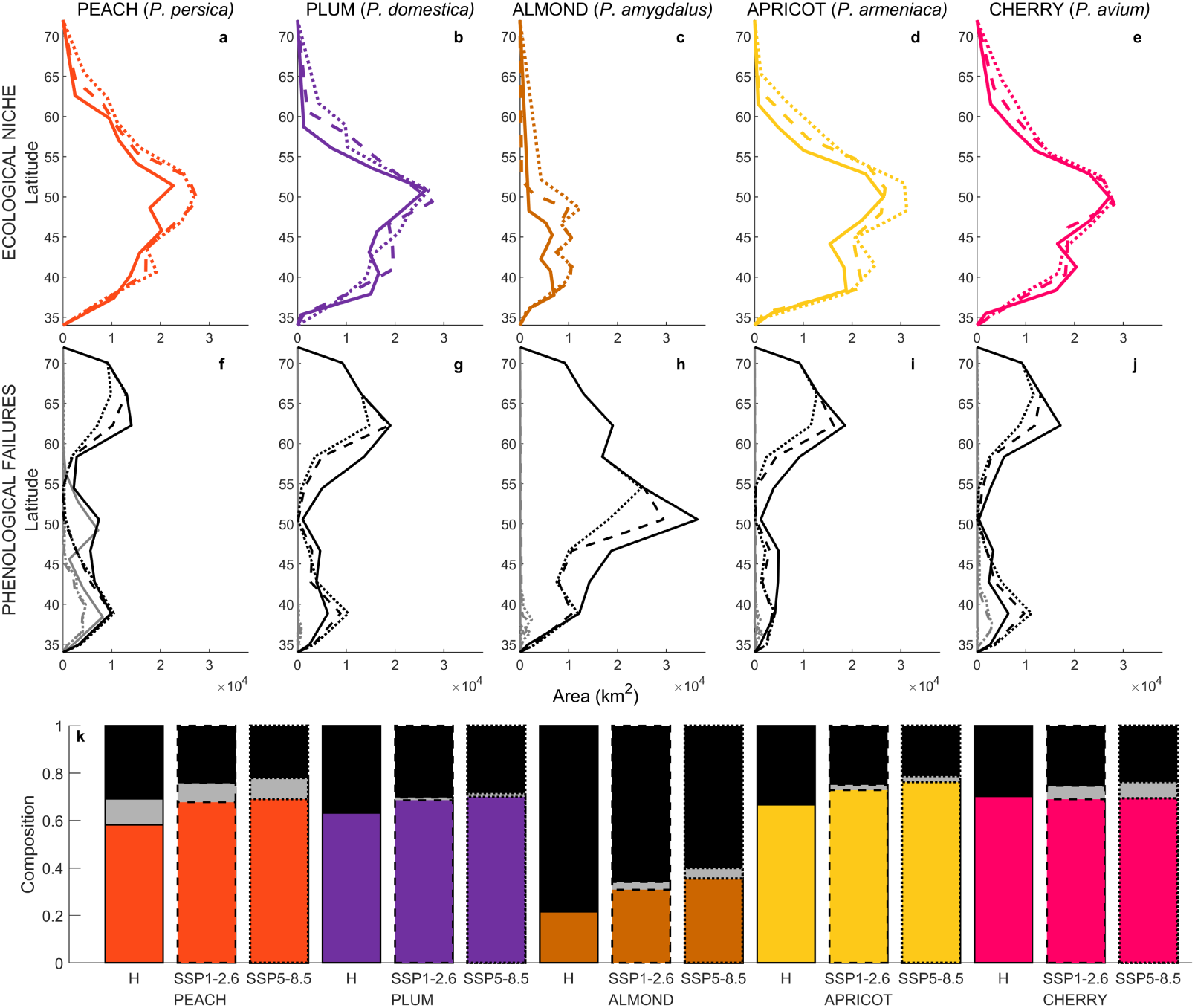
Latitudinal gradient of *Prunus* (a-e) ecological niche and (f-j) phenological failures (in gray chilling failures and in black forcing failures) in the historical period (1995-2014, solid line), at the middle of the century under SSP1-2.6 scenario (2041-2060, dashed line), and at the middle of the century under SSP5-8.5 scenario (2041-2060, dotted line). (k) Barplot represents the composition of the European territory between suitable areas (colored in orange for peach, purple for plum, brown for almond, yellow for apricot and magenta for cherry) in gray chilling failures and in black forcing failures, during the historical period (H, solid edge) and at the end of the century under the two scenarios (SSP1-2.6 scenario dashed bar edge, SSP5-8.5 scenario dotted bar edge).

